# Negative plant-microbiome feedback limits productivity in aquaponics

**DOI:** 10.1101/709162

**Authors:** Jessica A. Day, Anne E. Otwell, Christian Diener, Kourtney E. Tams, Brad Bebout, Angela M. Detweiler, Michael D. Lee, Madeline T. Scott, Wilson Ta, Monica Ha, Shienna A. Carreon, Kenny Tong, Abdirizak A. Ali, Sean M. Gibbons, Nitin S. Baliga

## Abstract

The demand for food will outpace productivity of conventional agriculture due to projected growth of the human population, concomitant with shrinkage of arable land, increasing scarcity of freshwater, and a rapidly changing climate. Efforts to increase conventional agricultural output come with significant environmental impacts stemming from deforestation and excessive use of chemicals, including soil salinization, erosion, and nutrient runoffs. While aquaponics has potential to sustainably supplement food production with minimal environmental impact, there is a need to better characterize the complex interplay between the various components (fish, plant, microbiome) of these systems to optimize scale up and productivity. For instance, much of our knowledge of beneficial and detrimental microbial communities vis-à-vis crop productivity comes from studies on plant-microbiome interactions in soil. Here, we investigated how the practice of continued transfer of microbial communities from pre-existing systems might promote or impede productivity of aquaponics. Specifically, we monitored plant growth phenotypes, water chemistry, and microbiome composition of rhizospheres, biofilters, and fish feces over 61-days of lettuce (*Lactuca sativa)* growth in aquaponic systems inoculated with bacteria that were either commercially sourced or originating from a pre-existing aquaponic system. Strikingly, *L. sativa* plant and root growth was significantly reduced across all replicates inoculated with the established microbiome. Further analyses revealed the reduced productivity was potentially a consequence of plant-specific pathogen enrichment, including *Pseudomonas*, through transfer of microbiomes from pre-existing systems – a phenomenon consistent with negative feedbacks in soil ecology. These findings underscore the need for diagnostic tools to monitor microbiome composition, detect negative feedbacks early, and minimize pathogen accumulation in aquaponic systems.

## Narrative

Sustainable food production has been on the rise in recent decades as traditional agricultural practices, which contribute to large-scale environmental degradation and enormous resource consumption, fall short of fulfilling the demands of our growing human population [1]. Aquaponics offers a sustainable alternative to traditional food production methods by combining hydroponic plant cultivation with aquaculture in a semi closed-loop system [2] that minimizes water and fertilizer use, increases agricultural efficiency [3], and does not require arable land. Central to the health of fish and plants in these systems are microorganisms, which drive many critical functions such as nitrogen cycling, plant growth promotion, disease resistance, and nutrient uptake; however, a deeper understanding of microbial community composition and function in aquatic agricultural systems is central to engineering and scaling-up efficient, sustainable food systems with low natural resource dependence [4]. While some work has been conducted on microbial communities in hydroponics [5] and aquaponics [4, 6], our knowledge of the microbial ecology of aquaponics is mainly grounded in soil-based agricultural research [7–9].

Until recently, interest among researchers and growers in aquaponic microbes has been focused on initiating nitrogen cycling and promoting plant growth via inoculation with plant growth promoting microbes (PGPMs). For this reason, one of two inoculation strategies are traditionally used to initiate cycling: 1) addition of commercially-derived nitrifying bacteria (*Nitrosomonas, Nitrobacter*, and *Nitrospira*) or 2) transfer of established bacteria from existing, healthy aquaponic systems. Despite the inclusion of PGPMs, a 2018 international survey found that 84.4% of aquaponic growers observed disease in their systems, of which 78.1% could not identify the causal agent [10]. Therefore, understanding the effect that microbial transfer has on plant production in aquaponics is crucial not only to establish best practices and increase commercial profitability by way of improving efficiency, but also to decrease loss due to disease. Of the growers who observed disease, a mere 6.2% used pesticides or biopesticides against plant pathogens and relied, instead, on non-curative actions, likely due to a lack of knowledge among aquaponic growers regarding plant pathogens and disease control [10]. Knowledge of key associations between microbial genera and plant productivity throughout early stages of system establishment could enable the development of diagnostic tools for monitoring microbiome composition, potentially aiding in early detection and prevention of system collapse.

Here, we examined how microbiome transfer from pre-existing systems might promote or impede plant productivity in aquaponics. We compared lettuce (*Lactuca sativa var. crispa*) growth in two distinct systems – those inoculated with a commercially-available microbial consortium (“commercial inoculum treatment” or “CIT”) and those inoculated with the biofilter media from an established, fully-cycled aquaponic system (“established inoculum treatment” or “EIT”).

Two sets of triplicate aquaponic systems were constructed using a 4-compartment design (Fig 1A). CIT systems were inoculated with Microbe-lift Nite-Out II PB commercial bacteria (Cape Coral, Florida, USA), marketed as containing key nitrifiers (*Nitrosomonas, Nitrobacter*, and *Nitrospira)*, whereas the established treatment consisted of systems inoculated with microbes from a previously established, healthy, highly-efficient *L. sativa* producing aquaponic system where nitrogen had been fully cycled for approximately 2 years. Microbiome samples were collected over a 61-day study period and analyzed from 3 compartments in each system: 1) plant roots in the grow bed, where PGPMs and plant pathogens are typically located, 2) biofilter media in the biofilter where nitrifiers typically carry out nitrification, and 3) fish feces from the solids filter, where anaerobic organisms and fish-specific pathogens can be found (Fig 1A). The sampling schedule was aligned with key chemical transformations during nitrification (“T0”, pre-cycling; “T1”, during cycling; “T2”, cycled), to capture associated microbial community shifts (Fig. 1A). Microbial gDNA was isolated from samples (Qiagen PowerSoil and PowerBiofilm kits; see supplement for methods), full-length 16S rRNA genes were amplified, and amplicons were sequenced with a MinION Nanopore Sequencer (see supplement for kits and methods).

**Figure 1.**
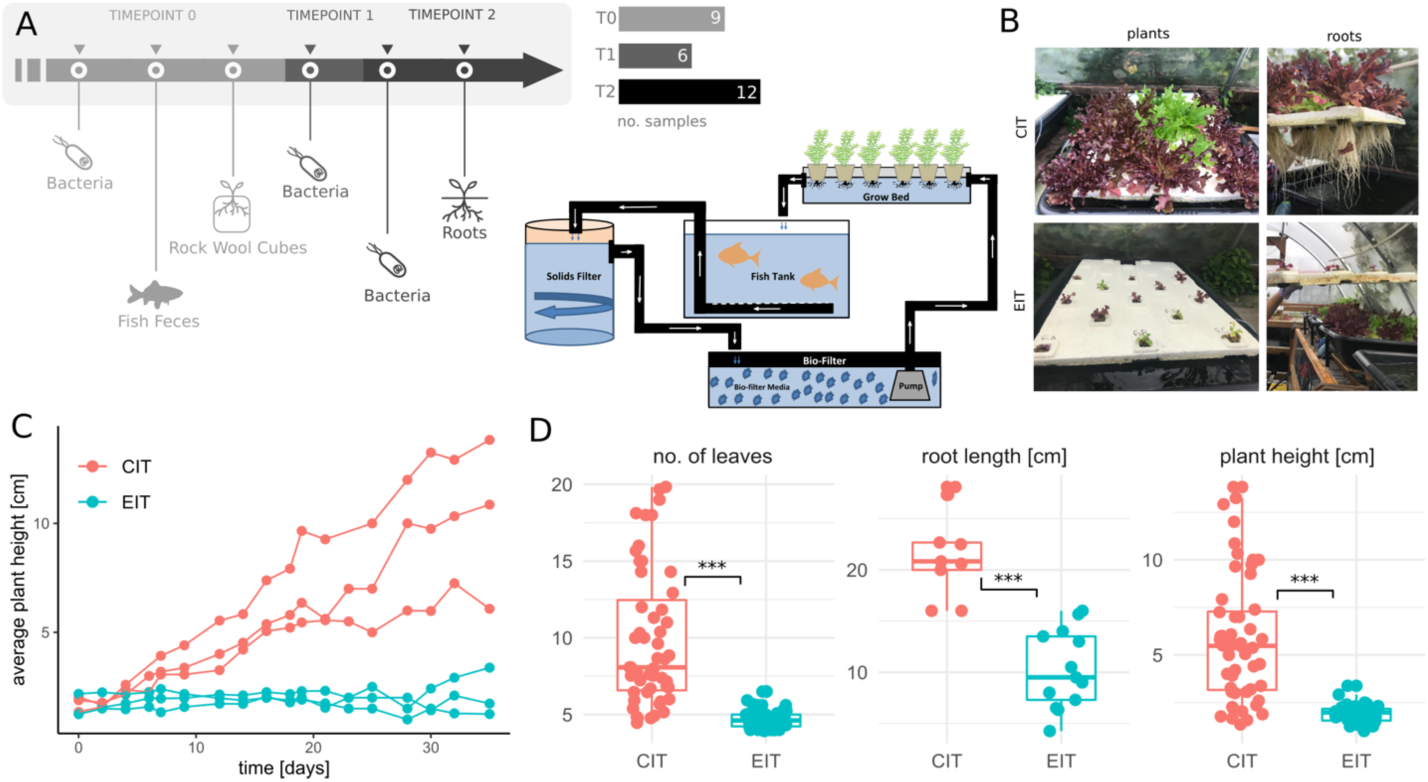
Aquaponic system design and plant phenotypes. (A) Aquaponics sampling timeline and system design. (B) Representative images of *L. sativa* plants and roots after a month of growth in systems with different inocula. (C) Plant growth over time. Each dot denotes the average plant height for a single aquaponic system taken at the indicated time point. Measures from the same tank are connected by lines. Gray line denotes growth in the prior aquaponics system that was used as the source for EIT. (D) Plant growth measures by inoculum. Each point denotes an average value measured in a single tank at a single time point (n = 104, 22, 105 for leaves, root length, and plant height, respectively). Stars denote significance under a Mann-Whitney rank-sum test (all p<0.001).

*L. sativa* growth (height, number of leaves, and root length) was significantly reduced in all EIT replicates compared to plant growth in CIT replicates (Fig 1B and C). Due to the consistency of physicochemical properties in all aquaponic systems across the study period (Fig. S1), it is unlikely the observed growth disparity can be explained by biogeochemical parameters. We investigated potential causes for the drastic reduction in plant growth in EIT replicates by exploring associations between microbial community composition and plant growth parameters.

We first investigated whether microbiome transfer affected establishment of microbes responsible for key nitrogen transformation processes. Previous studies of established aquaponic systems have found a ubiquitous presence of nitrifiers such as *Nitrosomonas, Nitrobacter*, and *Nitrospira*, albeit in low abundances, and these organisms are described as major drivers of plant growth [6, 11]. Our data revealed a very different scenario for systems in early establishment (Fig. S2). With the exception of fish feces, nitrifiers were not detected in microbial communities, which were dominated by Proteo- and Cyanobacteria (Fig. S3 and Fig. 2A, see supplement). Even though our protocol was validated to detect nitrifiers (Fig. S4), we also found no nitrifiers in the commercial inoculum, which was dominated by *Rhodanobacter*, a known denitrifying genus [12]. This suggests that the improved plant growth in CIT systems is independent of nitrifying bacteria. We also observed low nitrogen levels in our endpoint plant nutrient analyses (Table S2) and water chemistry (Fig. S1, S2), suggesting that nitrogen may have been limiting. This finding could explain the increased abundance of nitrogen fixers, such as *Rahnella*, and reduced abundance of nitrifying species (Fig. 2A). We hypothesize that in nitrogen-limited aquaponic systems, nitrogen fixing bacteria play an important role in supplementing the limited ammonia produced by fish by fixing atmospheric nitrogen and producing additional ammonia, which is a well-known nitrogen source for *L. sativa [13]*.

**Figure 2.**
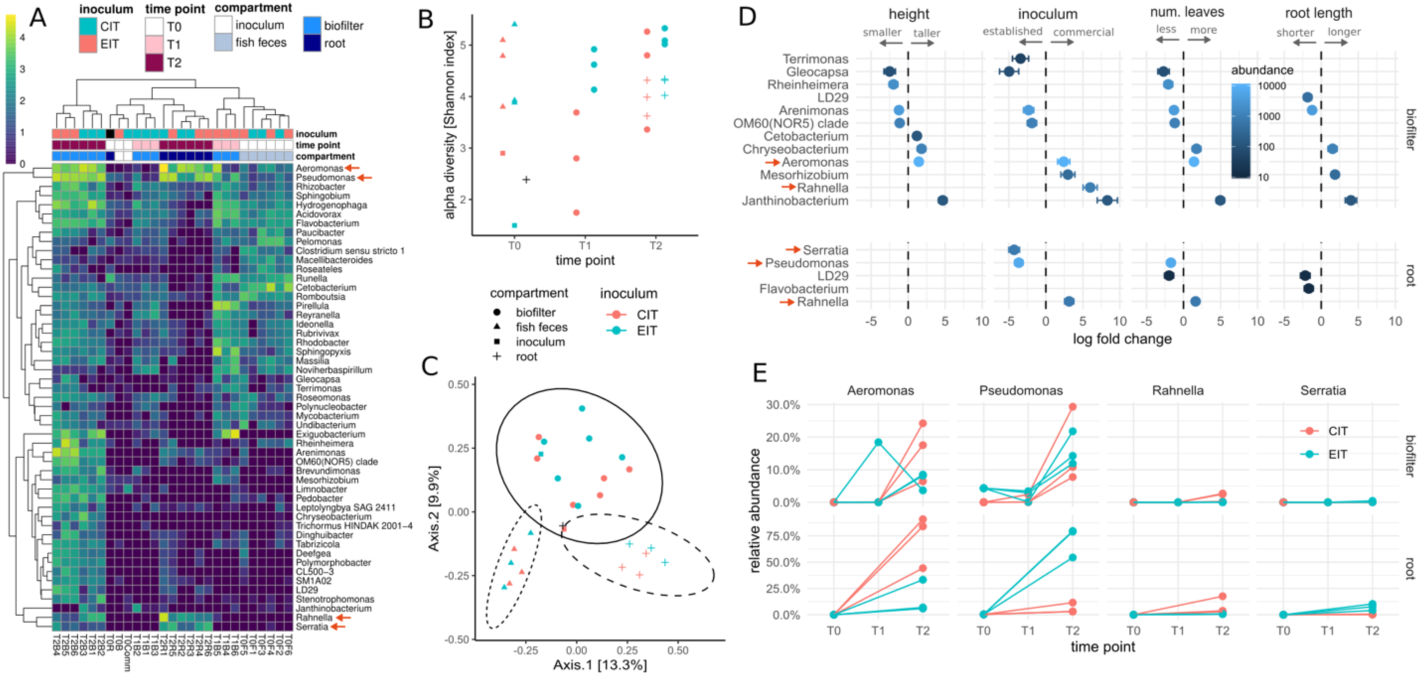
Sequencing of the complete 16S rRNA gene across the aquaponic system. (A) Abundances of ubiquitous bacterial genera across the samples (present in at least 2 samples at an abundance >300 reads). Colors of cells denote the normalized abundance on a base 10 log-scale. Sample names are composed of time point (e.g. T1), compartment (B = biofilter, F = fish feces, R = root, Comm = commercial inoculum) and tank number (1-6). Orange arrows denote genera of interest. Black fill color in A-C denotes initial root sample from rock wool cubes (not part of inoculation strategy). (B) Alpha diversity (Shannon index) over time. Colors denote initial source and shape compartment respectively. All samples were rarefied to 3,000 reads each. (C) PCoA plot of individual samples. Colors and shapes are the same as in B. All samples were rarefied to 3,000 reads each. Ellipses denote 95% confidence interval from Student t-distribution separating compartments. (D) Significant associations (FDR adjusted p<0.05) between bacterial genera and plant growth metrics or inoculum. Circles denote the association coefficient in the respective regression and error bars denote the standard error of the coefficient. Fill color denotes average abundance across all samples. Orange arrows mark genera of interest. (E) Time course of selected genera associated with plant growth and inoculum. Time point zero is shared between all samples and denotes initial inoculum for biofilter and initial plant microbial composition for roots. Only genera with more than 300 reads in at least 2 samples were considered in A-E (see supplement).

In examining whether microbiome transfer affected establishment of microbial communities in new systems, we found alpha-diversity increased with time in all compartments and achieved similar values for CIT and EIT tanks in biofilters and roots at T2 (Fig. 2B). Aquaponic compartments each had distinct microbial compositions (Fig. 2B). A total of 44% of variation in beta-diversity was explained by a combination of compartment (25%), inoculum (11%), and an interaction term of the two (8%; all PERMANOVA p values < 0.02). We also found that samples clustered by time point (Fig. 2C). Conversely, prior studies have found that the microbial composition of different compartments in established systems, with the exception of fish feces, are quite similar [6].

Given the differences in microbial composition between EIT and CIT systems, we hypothesized there may be a negative effect of microbial transfer on plant growth, which acts independent of nitrogen concentration and cycling timeline. Thus, we investigated whether the difference in plant growth could be explained by an enrichment in potentially growth-inhibiting bacteria from the established system inoculum. This microbiome-inhibition hypothesis is similar to negative plant-soil feedback (NPSF), which has long been studied in soils [14, 15]. NPSF is characterized by enrichment of species-specific plant pathogens limiting plant productivity in successive generations grown in the same soil. This self-inhibitory process promotes plant community diversity by imposing greater mortality on established species and allowing sub-dominant species to thrive in their place [14, 15]. Because we hypothesize that a similar phenomenon might be at play in a soil-free environment, we will henceforth use the term “negative plant-microbiome feedback” (NPMF).

In order to identify biotic mechanisms of plant-microbiome feedbacks, we examined whether there were distinct bacterial genera in the plant roots that were associated with inoculum source and plant growth phenotype. Association tests between bacterial genus-level abundances and plant growth measures revealed that 5 genera in the roots were significantly associated with plant growth (FDR corrected p<0.05), of which *Pseudomonas, Rahnella* and *Serratia* were the most abundant (Fig. 2D, see supplement for methods). *Pseudomonas* and *Serratia* were higher in the EIT systems and were associated with diminished plant growth, whereas *Rahnella* was higher in CIT systems and was associated with improved plant growth (Fig. 2D). Although these genera were all associated with plant growth metrics, only *Pseudomonas* was 1) previously described as a plant pathogen [16–18] and 2) found in both initial inoculum types. Across time points, we observed a relatively uniform accumulation of *Pseudomonas* in biofilters, with all biofilter microbiomes consisting of 5-30% *Pseudomonas* (Fig. 2E). However, detection of *Pseudomonas* in the rhizosphere was highly dependent on inoculum type. EIT rhizosphere microbiomes were dominated by *Pseudomonas* (with 50-80% *Pseudomonas*, DESeq2 FDR-corrected p<0.05), whereas CIT rhizosphere microbiomes contained less than 20% *Pseudomonas*. Instead, CIT rhizospheres were enriched for *Aeromonas* and *Rahnella*, which were associated with improved plant growth (Fig. 2D). In summary, these findings suggest that *Pseudomonas* proliferates within biofilters across all systems, but only dominates the *L. sativa* rhizosphere in systems inoculated with microbes from an established aquaponic system.

While future work with larger sample sizes will be necessary to characterize putatively pathogenic strains, establish causality, and verify our results, our findings suggest that negative plant-microbiome feedback may be a major determinant of agricultural performance in early and nitrogen-limited aquaponic systems. This extends an important lesson from soil ecology to aquatic agriculture. While NPSF can be beneficial in natural environments by means of increasing biodiversity [19, 20], it can be detrimental in agricultural systems when cultivating a single crop species without incorporating crop rotation [21]. Furthermore, the decreased yield (∼36-fold decrease in plant biomass; Table S1) observed in this study could be financially devastating to aquaponic farmers [22]. Future studies should identify the effect of crop rotation and/or multi-crop aquaponic systems on plant growth and pathogen enrichment. Because our analyses did not detect nitrifying bacteria in the original commercial inoculum, the most cost-effective inoculation strategy could be allowing systems to naturally accumulate microbes from the ambient environment, although additional studies without added microbes would be needed to confirm this hypothesis. If this is the case, it may be more important to combat the steady accumulation of pathogens in monoculture systems, possibly with biocontrol agents [10, 23], rather than to seed the system with beneficial microbes. Moving forward, metagenomic analyses will provide deeper insights into the specific species/strains and functional genes involved in plant-microbiome feedbacks [24] and could lead to more targeted strategies for early detection of pathogens and pathogen suppression.

## Acknowledgements

This research was possible due to the strategic collaborations fostered by Claudia Ludwig, as well as the generous support and resources provided by Claudia Ludwig, Ray Williams and the Black Farmers Collective, Jeff King and the Microsoft Giving Campaign, Fred Hutchinson Cancer Research Center, Seattle Youth Employment Program, CrowdRise donors, the National Science Foundation (NSF MCB-1616955, MCB-1518261, DBI-1565166, MCB-1330912), a Washington Research Foundation Distinguished Investigator Award (supporting CD and SMG), and a Scientific Innovation Fund grant from the NASA Office of the Chief Scientist to Brad M. Bebout.

