## Supplementary Information for "Negative plant-microbiome feedback limits productivity in aquaponics"

**Manuscript title: "Negative plant-microbiome feedback limits productivity in aquaponics"**

Document Summary

This supplemental document includes content, figures, and tables regarding experimental system design, system monitoring and sampling methods, sample preparation and sequencing, and data analysis and validation that supports the manuscript "Negative plant-microbiome feedback limits productivity in aquaponics". Data and software availability information is included at the end of this document.

*Aquaponic system design*

Two sets of triplicate aquaponic systems, treated with either commercial or established inoculum, were constructed using a 4-compartment design (Fig 1A) in an open-air greenhouse in Seattle, Washington. These treatments will henceforth be labeled CIT (commercial inoculum treatment) and EIT (established inoculum treatment). Each system housed one aquaculture tank connected to one hydroponic unit utilizing the deep water culture method. These units were separated by a biofilter, which housed approximately 60L of K3 filter media (bioballs), as well as an 18 L solids filter. The experiment commenced when two adult Red Nile Tilapia (*Oreochromis niloticus*; the sum of their lengths being approximately 44cm), were added to each system. On the same day, the 6 systems were divided into two sets of triplicate systems and inoculated with either Microbe-Lift commercial bacteria (Cape Coral, Florida, USA; tanks 1-3; "CIT") or established microbes from an existing aquaponic system (tanks 4-6; "EIT"). Four weeks later, 13 Oak Leaf Blend lettuce (*Lactuca sativa var. crispa*) seedlings, which had been sprouted in rockwool cubes outside the systems 3 weeks earlier, were added to the floating trays (Beaver Plastics 28-hole 2ft x 4ft Lettuce Raft) in each system and allowed to grow 35 days.

*Animal Care*

Prior on-site research conducted at the Institute for Systems Biology under IACUC protocol number NB.01a.16 ("Educational Aquaponics"; OLAW #A4355-01 and AAALAC #001363) provided guidelines that were closely followed in this off-site study. Water chemistry, space, and fish food were all determined and maintained to optimize living conditions for all Nile Tilapia (*Oreochromis niloticus*) used in this study, and fish health and behavior were observed and recorded regularly. Fish were fed Stage 3 Intermediate AquaNourish Omnivorous Fish Feed pellets (Star Milling Co., Perris, California, USA; 37% crude protein, 10% crude fat, 2.2% crude fiber, 11% ash) every other day in an increasing amount (between 10-40 pellets per fish) during nitrogen cycling, so as not to produce excess, toxic ammonia at initial stages of cycling. Each approximately 203 L system housed two fish with a total length of 44 inches. Throughout this study, 5 fish died unexpectedly due to unknown causes in tanks 1, 2, 3, and 6. Water chemistry was ruled out as a cause, as no associations between water chemistry fluctuations and subsequent fish deaths were found. One possible cause may be infection by a fish pathogen such as *Saprolegnia* or *Aphanomyces*. Although 16S rRNA analysis is not capable of detecting these fungal pathogens, *Pseudomonas* and *Aeromonas* were increasingly enriched throughout the study. Both of these genera are known to include species which can mitigate oomycete diseases in aquaculture by acting as an antipathogen agent [1].

*Inoculation*

Systems in the CIT were inoculated with Microbe-lift commercial bacteria, marketed as containing *Nitrosomonas, Nitrobacter,* and *Nitrospira*, whereas EITs were inoculated with biofilter media from an established aquaponic system which had previously been a highly efficient producer of *L. sativa* for approximately 2 years. To inoculate these systems, 10 bioballs (representing approximately 105.8 ng/mL bacteria DNA) from the existing aquaponic system were collected and transferred to the biofilter of each new EIT system. On the same day, 30 mL of commercial bacteria (company recommendation) was added to the CIT biofilters.

*Water chemistry maintenance*

Throughout the study period, water chemistry (pH, temperature, ammonia, nitrite, nitrate, chlorine, hardness, and alkalinity) and plant growth metrics (number of leaves, plant height, root length, and presence of disease or discoloration) were collected 3-4 times per week (Fig S1). Water chemistry data was collected using test strips (Tetra EasyStrips 6-in-1 Aquarium Test Strips and Tetra EasyStrips Ammonia Aquarium Test Strips) and a multi-parameter probe (Hanna Instruments® GroLine Waterproof Portable pH/EC/TDS Meter). To account for evaporation and plant use, an average of 126.5L of aerated, dechlorinated tap water was added to each tank throughout the experimental period. Approximately 45g of Instant Ocean® Sea Salt (Blacksburg, VA) was added to each tank 5 days into the experiment to reach a conductivity of approximately 900 uS/cm.

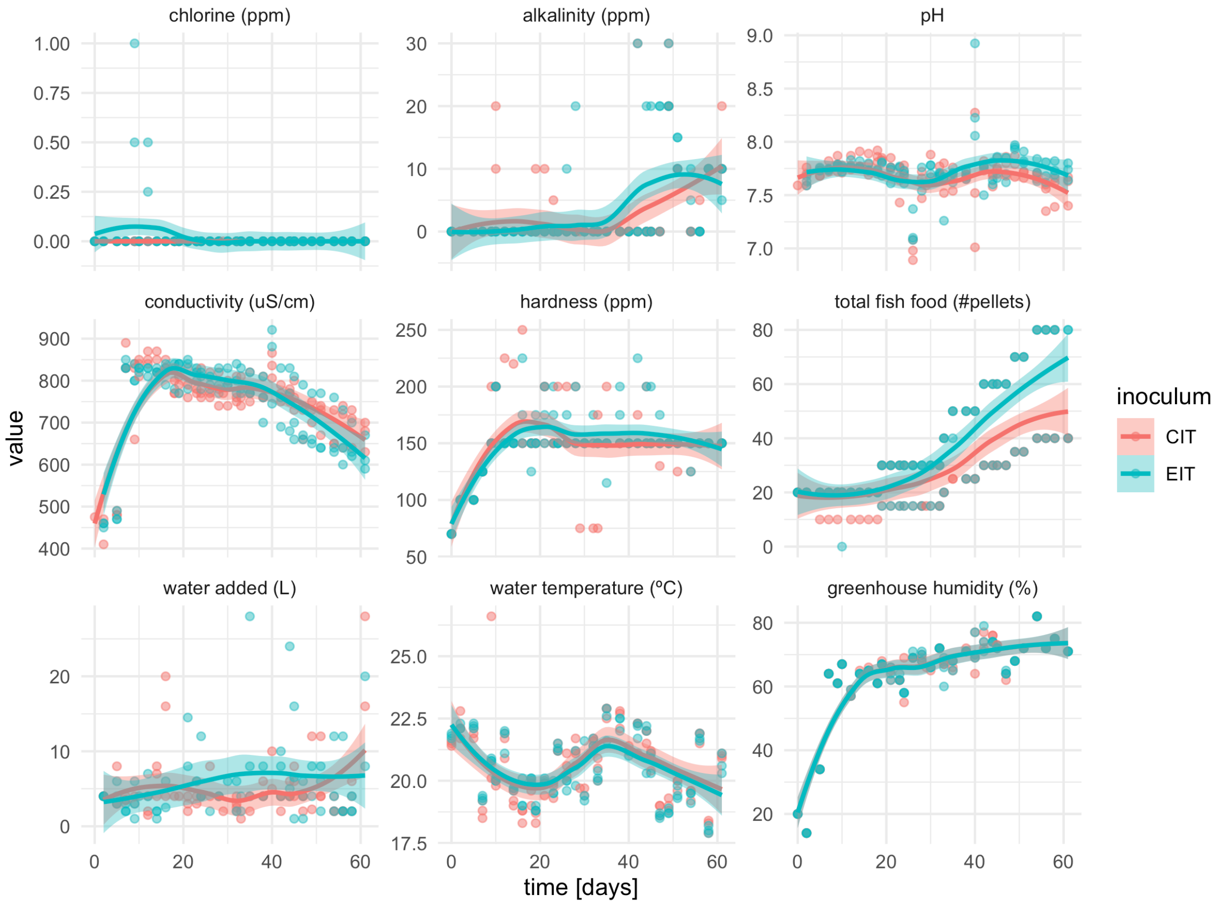


Figure S1. Water chemistry, environmental parameters, and system inputs measured throughout the study period. Lines denote LOESS smoothed curves for each inoculum and bands denote 95% confidence intervals of the smoothing function.

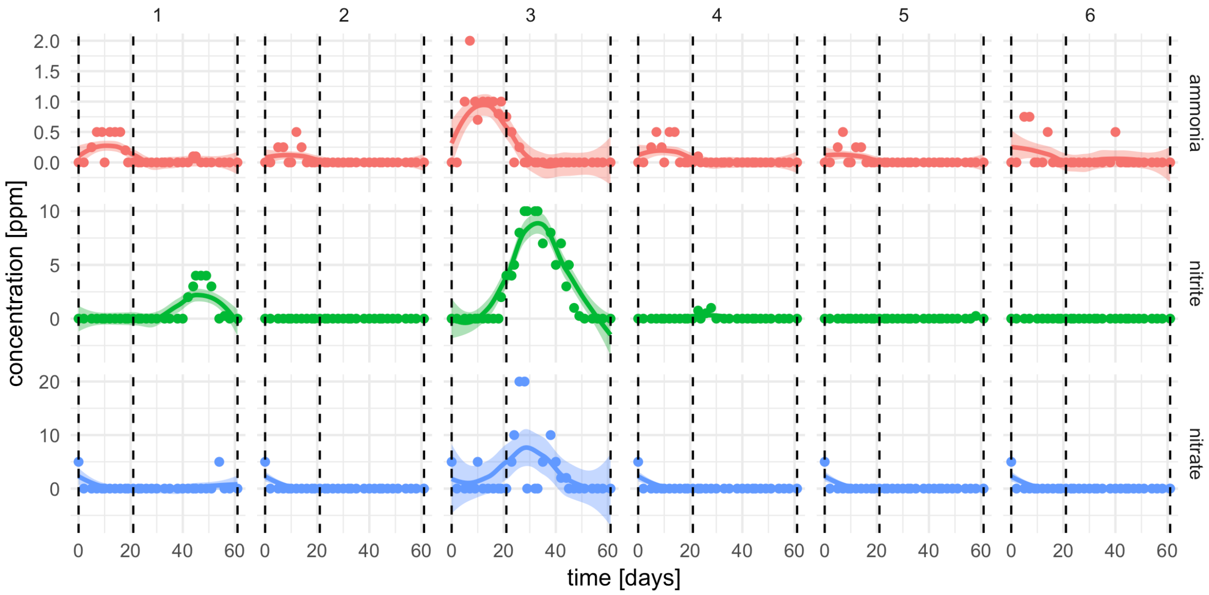


Figure S2. Measures of nitrogen cycling. Colored lines denote smooth fit from a LASSO regression and filled areas denote 95% confidence intervals. Numbers of the panels denote tanks. Tanks 1-3 are CIT whereas tanks 4-6 are EIT. Dashed lines denote the three time points used for microbiome sampling.

*Sample collection*

Microbiome samples were strategically collected throughout the 61-day study period and analyzed from 3 compartments of each system: 1) plant roots in the grow bed, 2) biofilter media in the biofilter, and 3) fish feces from the solids filter (Fig 1A). Sampling time points were reflective of key chemical species transformations occurring during nitrification, and therefore supposed microbial community shifts, across the study period (pre-cycling, during cycling, and post-cycling; Fig 1A). The experiment ended 61 days after microbial inoculation and plants were harvested. At the end of the experiment, wet mass and dry mass (biomass) of plants was collected (Table S1). To obtain total biomass, leaves were placed in an oven at 60C for 72 hours and weighed immediately after being removed. Nutrient analysis of dried leaf samples was performed by the

University of Missouri Soil and Plant Testing Lab (Table S2).

Table S1: Final wet mass and dry mass (biomass) of *L. sativa* leaves in all replicates.

| ***Total mass (g)*** | ***CIT System Replicates*** | | | ***EIT System Replicates*** | | |
|---|---|---|---|---|---|---|
| | ***1*** | ***2*** | ***3*** | ***4*** | ***5*** | ***6*** |
| Wet | 238.191 | 52.221 | 329.201 | 1.819 | 1.702 | 8.301 |
| Dry | 18.392 | 4.719 | 24.529 | 0.339 | 0.268 | 0.722 |

Table S2: Amount of 9 key nutrients found in *L. sativa* leaves at the end of the 61-day study period. *Low concentrations compared to *L. sativa* soil-grown sufficiency range. **High concentrations compared to *L. sativa* soil-grown sufficiency range.

| ***Nutrient*** | ***CIT System Replicates*** | | | ***EIT System Replicates*** | | |
|---|---|---|---|---|---|---|
| | ***1*** | ***2*** | ***3*** | ***4*** | ***5*** | ***6*** |
| *Nitrogen (N) %* | 2.37* | 1.93* | 2.13* | 1.35* | 1.90* | 3.20* |
| *Phosphorus (P) %* | 0.37* | 0.23* | 0.36* | 0.09* | 0.08* | 0.30* |
| *Potassium (K) %* | 4.12* | 4.47* | 4.29* | 1.74* | 1.55* | 4.72* |
| *Calcium (Ca) %* | 1.35 | 1.32 | 1.62 | 1.38 | 1.28 | 1.39 |
| *Magnesium (Mg) %* | 0.54** | 0.49** | 0.65** | 0.44** | 0.47** | 0.47** |
| *Iron (Fe) ppm* | 229** | 279** | 106 | 412** | 243** | 251** |
| *Manganese (Mn) ppm* | 22 | 27 | 12 | 38 | 62 | 42 |
| *Copper (Cu) ppm* | 6 | 7 | 3 | 23 | 7 | 7 |
| *Zinc (Zn) ppm* | 103 | 180 | 160 | 76 | 64 | 131 |

*Sample processing*

***Roots:*** Time point 0 for the roots was taken from rockwool cubes that seedlings were growing in prior to transfer to the aquaponic systems. 9 rockwool cubes were collected from the grow tray and all liquid was squeezed from the rockwool into a 50 mL falcon tube. Tubes were centrifuged at 4,000g for 10 min, supernatant was discarded, and pellets were flash frozen in liquid nitrogen and stored at -80C until DNA extraction. The final time point for the roots was taken at the time of system disassembly (and plant harvest). Roots were clipped at the base of all plants in each system and collected in 50 mL falcon tubes. PBS was added to the tube and samples were sonicated at low frequency (intensity 1) for 5 minutes total (five 30 second bursts, each followed by a 30 second rest period) in order to remove cells from roots. Following sonication, roots were removed from the tube and samples were centrifuged at 4,000g for 10 min, supernatant was removed, and pellets were flash frozen in liquid nitrogen and stored at -80C until DNA extraction.

***Fish feces:*** Fish feces were collected from the bottom of the aquaponic solids filter using a 25 mL serological pipette. Samples were centrifuged at 4,000g for 10 minutes to pellet the feces samples, supernatant was discarded, and pellets were flash frozen in liquid nitrogen and stored at -80C until DNA extraction.

***Biofilter:*** Ten bioballs were collected from the established aquaponics biofilter using sterile forceps and stored in PBS. A single bioball was vortexed at maximum speed for 2 minutes in a 50 mL falcon tube and then sonicated at low frequency (intensity 1) for 5 minutes total (five 30 second bursts, each followed by a 30 second rest period). Following sonication, the bioball was removed from the tube and another bioball was added in, vortexed, and sonicated. This process was repeated until cells from 10 bioballs were collected for each sample. Cells from the 10 bioballs were then collected as a single pellet through centrifugation at 4,000g for 10 minutes, supernatant was discarded, and pellets were flash frozen in liquid nitrogen and stored at -80C until DNA extraction.

*DNA extraction, 16S rRNA sequence analysis*

Microbial gDNA was isolated from samples using 2 similar kits marketed specifically for bacterial isolation from agricultural environments (Samples T0F1, T0F2, T0F3, T0F4, T0F6, and T0B with Qiagen PowerSoil kit and all others with PowerBiofilm kit), 16S rRNA genes were amplified using universal primers suggested by MinION - 27F 5'–AGAGTTTGATCMTGGCTCAG and 1492R 5'–CGGTTACCTTGTTACGACTT [2], and a MinION Nanopore Sequencer was used for sequence analysis.

*Sequencing*

Amplicons were aliquoted to a starting concentration of 1ug and were further processed and sequenced according to ONT's 1D Native barcoding genomic DNA (with EXP-NBD103 and SQK-LSK108) protocol (v. NBE_9006_v103_revN_21Dec2016), beginning with an AMPure XP bead purification step, and proceeding to the end-repair/dA-tailing, barcoding ligation, and adapter ligation steps.

*Analysis*

Basecalling for the raw MinION files was performed by Albacore (v2.0.2) to yield the corresponding FASTQ files. Reads were processed using "filterAndTrim" methods from DADA2 [3]. The first 10bp of the 5' end were trimmed from all reads to as they generally showed lower qualities. Raw reads were also trimmed to a maximum length of 1.5kbp (the expected length of the 16S gene). Reads with more than 200 expected errors under Illumina error model (based on [4]) or more than 2 ambiguous base calls ("N" bases) were removed from the analysis (~70% of all reads passed these filters). The filtered reads were then aligned to the complete SILVA 16S database using minimap2 with the Oxford Nanopore preset allowing up to 100 alternative alignments per read [5, 6].

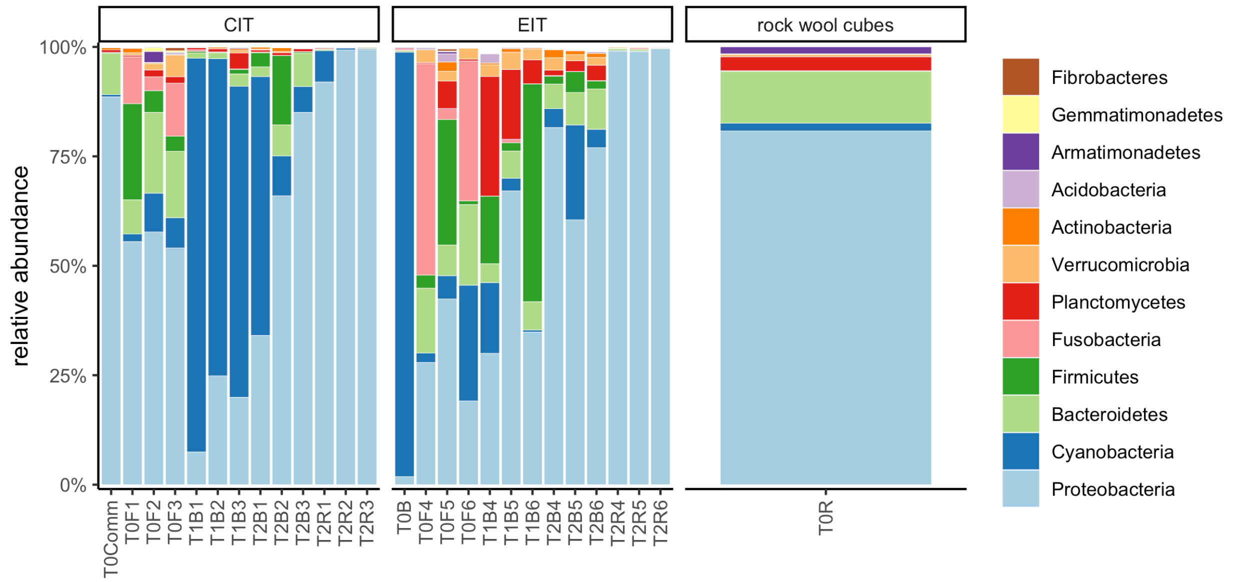


Figure S3. Relative abundances for the 12 most abundant taxa in the data set. Sample names are composed of time point (e.g. T1), compartment (B = biofilter, F = fish feces, R = root, Comm = commercial inoculum) and tank number (1-6). Panels denote inoculum as used in Figure 1 of the main text.

*Validation and mapping improvement for high error rates*

We observed acceptable read qualities with a median of 15 corresponding to an average error rate of 3% under the Illumina model. However, quality measures are based on a predictive model and the nominal error rate is usually above 10% [4]. In order to evaluate the accuracy of nanopore sequencing on the full length 16S gene we used a small validation data set consisting of two biological replicates of a long-running established aquaponic system sequenced with the same nanopore protocol as well as V4-V5 Illumina amplicon sequencing. V4-V5 sequencing data (515F 5'–GTGYCAGCMGCCGCGGTAA, 926R 5'–CCGYCAATTYMTTTRAGTTT; [7] were analyzed using the DADA2 pipeline and used as the ground truth. Due to the high error rate of nanopore sequencing we expected a lot of spurious assignments when mapping to a 16S reference database such as SILVA. In order to limit those spurious matches we employed an Expectation-

Maximization (EM) algorithm similar to what is used in kallisto, but without correction for gene length due to the fixed length of the 16S gene [8] (implementation available at https://github.com/Gibbons-Lab/mbtools). Applying the EM algorithm led to a reduction of unique mappings for low abundance cutoffs compared to a “naive” counting algorithm that just uses the highest scoring match (Figure S4A). We observed good agreement of estimated abundances between Illumina and Nanopore sequencing down to the genus level if the taxon was observed in both sequencing technologies (Figure S3B, $R^2$ between 0.5-0.8, Spearman rho between 0.73 - 0.85, all ANOVA $p<10^{-6}$). However, we also observed many spurious mappings only present in the nanopore data, with generally low abundances (Figure S4B-C). We found that using an abundance cutoff of read 300 counts removed >96% of all spurious mappings across all taxa ranks and >97% of all spurious mappings on the genus rank. Thus we employed this abundance cutoff throughout the manuscript. The final abundances as estimated by the EM algorithm were then annotated by the SILVA taxonomy down to the strain level where available and collapsed on the genus level [9]. We also verified whether the nanopore 16S primers were capable of identifying the same nitrifiers that were amplified by the V4-V5 primers used in the Illumina run. We found that all known nitrifying taxa identified by Illumina sequencing were also found in the Nanopore data (Figure S4D).

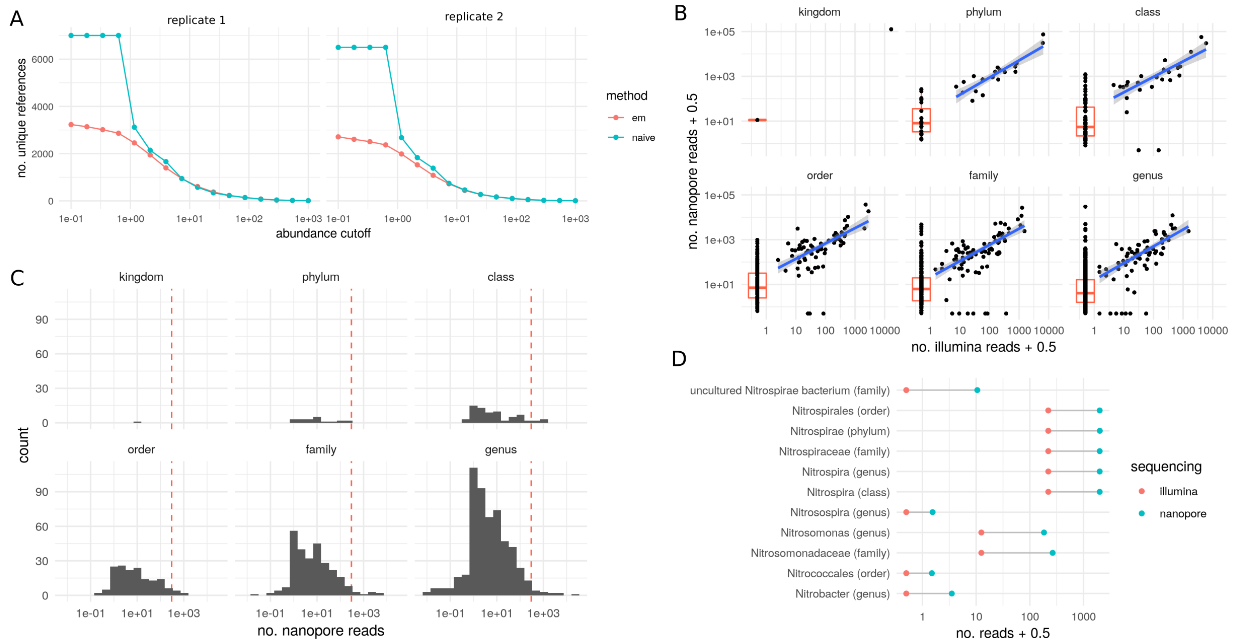


Figure S4. Validation of nanopore sequencing on a set of aquaponics biofilter samples that were sequenced using Illumina and Nanopore technologies (2 replicates each). (A) In low abundances, the Expectation-Maximization (em) algorithm identifies fewer unique references than the "naive" strategy of just selecting the highest scoring read. Each dot denotes the number of unique 16S sequences in the SILVA database that pass the abundance cutoff. (B) Abundances across sequencing protocols. Each dot denotes the abundance of a single taxon at the indicated taxonomic rank. Blue lines denote a linear regression for the taxa found in both sequencing technologies and gray errors denote the 95% confidence interval of the regression. The red boxplot summarizes the distribution of spurious mappings in the nanopore data (absent in the Illumina data). (C) Distribution of false positive mappings (mappings not observed in the Illumina data) in nanopore sequencing. The dashed red line denotes the used abundance cutoff that removed >96% of those spurious mappings. (D) Abundances of nitrifying taxa in both sequencing protocols. Dots denote the sum in the two replicates. Abundances smaller than one denote taxa not detected in Illumina sequencing.

*Association testing*

Read abundances across samples were normalized using the DESeq2 “poscounts” normalization strategy (modified robust centered log ratios) [10]. Association tests were performed using DESeq2 with a prior filtering step to remove bacterial genera with either very low average abundances or low prevalence across samples (mean abundance >10 across all samples and present in at least 2 samples). This more permissive filter was used to allow for effective shrinkage for DESeq2. Finally, we only considered significant tests for those genera which showed abundances larger than the default cutoff of 300 reads in at least two samples. Tests were performed for each of the 3 environments (biofilters, roots and fish feces) separately. False discovery rate was controlled by the Benjamini-Hochberg method [11]. No associations in the fish feces passed an FDR cutoff of 0.05.

*Data and software availability*

R notebooks and scripts to reproduce the analysis starting from the raw FASTQ files are provided at https://github.com/gibbons-lab/aquaponics. The analysis of the validation samples can be found at https://github.com/gibbons-lab/nanopore_vs_illumina. More complex algorithms such as the Expectation-Maximization algorithm are available in a dedicated R package along with documentation at https://github.com/gibbons-lab/mbtools. Raw sequencing data has been deposited in NCBI Sequence Read Archive (SRA) under the project IDs PRJNAXXXXXXX and PRJNAXXXXXX (validation data). **Data will be uploaded prior to publication.**

Supplemental References


1. Liu Y, Rzeszutek E, van der Voort M, Wu C-H, Thoen E, Skaar I, et al. Diversity of Aquatic Pseudomonas Species and Their Activity against the Fish Pathogenic Oomycete

Saprolegnia. *PLoS One* 2015; **10**: e0136241.

2. Stackebrandt E, Goodfellow M. Nucleic acid techniques in bacterial systematics. 1991. John Wiley & Son Ltd.
3. Callahan BJ, McMurdie PJ, Rosen MJ, Han AW, Johnson AJA, Holmes SP. DADA2: High-resolution sample inference from Illumina amplicon data. *Nat Methods* 2016; **13**: 581–583.
4. Teng H, Cao MD, Hall MB, Duarte T, Wang S, Coin LJM. Chiron: translating nanopore raw signal directly into nucleotide sequence using deep learning. *Gigascience* 2018; **7**.
5. Li H. Minimap2: pairwise alignment for nucleotide sequences. *Bioinformatics* 2018; **34**: 3094–3100.
6. Quast C, Pruesse E, Yilmaz P, Gerken J, Schweer T, Yarza P, et al. The SILVA ribosomal RNA gene database project: improved data processing and web-based tools. *Nucleic Acids Research* . 2012. , **41**: D590–D596
7. Parada AE, Needham DM, Fuhrman JA. Every base matters: assessing small subunit rRNA primers for marine microbiomes with mock communities, time series and global field samples. *Environ Microbiol* 2016; **18**: 1403–1414.
8. Bray NL, Pimentel H, Melsted P, Pachter L. Near-optimal probabilistic RNA-seq quantification. *Nat Biotechnol* 2016; **34**: 525–527.
9. Yilmaz P, Parfrey LW, Yarza P, Gerken J, Pruesse E, Quast C, et al. The SILVA and 'All-species Living Tree Project (LTP)' taxonomic frameworks. *Nucleic Acids Research* . 2014. , **42**: D643–D648
10. Love MI, Huber W, Anders S. Moderated estimation of fold change and dispersion for RNA-seq data with DESeq2.
11. Benjamini Y, Hochberg Y. Controlling the False Discovery Rate: A Practical and Powerful Approach to Multiple Testing. *Journal of the Royal Statistical Society: Series B (Methodological)* . 1995. , **57**: 289–300
